# Phenotyping the hidden half: Combining UAV phenotyping and machine learning to predict barley root traits in the field

**DOI:** 10.1101/2024.12.02.626299

**Authors:** Samir Alahmad, Daniel Smith, Christina Katsikis, Zachary Aldiss, Sarah V. Meer, Lotus Meijer, Karine Chenu, Scott Chapman, Andries B. Potgieter, Anton Wasson, Silvina Baraibar, Jayfred Godoy, David Moody, Hannah Robinson, Lee T. Hickey

**Affiliations:** Queensland Alliance for Agriculture and Food Innovation (QAAFI), The University of Queensland (UQ), Brisbane, QLD, 4072, Australia; School of Agriculture and Sustainability, UQ, St Lucia, QLD, 4072, Australia; UQ, QAAFI, Centre for Crop Science, Gatton, QLD, Australia; Agriculture & Food, Commonwealth Scientific and Industrial Research Organisation, Canberra, ACT, 2601, Australia; InterGrain Pty Ltd, Perth, WA 6163, Australia; Department of Plant Breeding, Hochschule Geisenheim University, 65366 Geisenheim, Germany

**Keywords:** Crop improvement, Haplotype mapping, LGEBV, Machine learning, Partial least square, Prediction, Random forest, Root phenotyping, UAV phenotyping, Vegetation indexes

## Abstract

Improving crop root systems for enhanced adaptation and productivity remains challenging due to limitations in scalable non-destructive phenotyping approaches, inconsistent translation of root phenotypes from controlled environment to the field, and a lack of understanding of the genetic controls. This study serves as a proof of concept, evaluating a panel of Australian barley breeding lines and cultivars (*Hordeum vulgare* L) in two field experiments. Integrated ground-based root and shoot phenotyping was performed at key growth stages. UAV-captured vegetation indices (VIs) were explored for their potential to predict root distribution and above-ground biomass. Machine learning models, trained on a subset of 20 diverse lines, with the most accurate model applied to predict traits across a broader panel of 395 lines. Unlike previous studies focusing on above-ground traits or indirect proxies, this research directly predicts root traits in field conditions using VIs, machine learning and root phenotyping. Root trait predictions for the broader panel enabled genomic analysis using a haplotype-based approach, identifying key genetic drivers, including *EGT1* and *EGT2* which regulate root gravitropism. This approach offers the potential to advance root research across various crops and integrate root traits into breeding programs, fostering the development of varieties adapted to future environments.

**Highlight:** Integrating UAV phenotyping and machine learning can be used to predict RSA traits non-destructively and offers a new approach to support root research and crop improvement.

## Introduction

Optimisation of root system architecture (RSA) has been flagged as a key breeding target to develop climate-resilient crops for the future (Ober *et al*., 2021). RSA refers to the spatial and temporal distribution of roots, which influences the uptake of water and nutrient resources in the soil to sustain and maximise crop performance (Koevoets *et al*., 2016). However, the value of different root architectures is highly context-dependent and driven by the environmental and management conditions experienced by the crop (Voss-Fels *et al*., 2018; Shazadi *et al*., 2024). For example, root proliferation in superficial soil layers could be advantageous in environments that experience sporadic rainfall and also facilitate access to nutrients, including phosphorous (Henry *et al*., 2010; Lynch, 2019; van der Bom *et al*., 2023). On the other hand, deeper root systems have the potential to enhance access to stored moisture to improve yield under terminal drought conditions (Rich *et al*., 2016; Alahmad *et al*., 2019). For instance, modelling studies suggest wheat yield could increase by 55 kg/ha for each additional millimetre of water extracted during grain filling. (Manschadi *et al*., 2006; Christopher *et al*., 2013). Customising crop root systems for specific agroecological contexts has the potential to achieve enhanced water and nutrient efficiency, sustainability and profitability (Ober *et al*., 2021).

Historically, RSA traits were overlooked in crop improvement programs, where breeders have focussed on above-ground traits, including flowering time and height (Den Herder *et al*., 2010; Borrell *et al*., 2014; Christopher *et al*., 2016; Robinson *et al*., 2018; Ober *et al*., 2021). The primary reason for this has been the difficulty of directly measuring RSA traits in the field. While innovations in field-based root phenotyping methods, including “mini-rhizotrons”, “shovelomics” and “core break” methods (Trachsel *et al*., 2011; Wasson *et al*., 2014; York *et al*., 2018) have provided advances and offer valuable insights, they are often constrained by population size and labour intensity (Trachsel *et al*., 2011; Wasson *et al*., 2014; York *et al*., 2018).

High-throughput, cost-effective techniques under controlled conditions, such as ‘paper pouch’ (Hund *et al*., 2009), ‘clear pot’ (Richard *et al*., 2015) and rhizobox methods (Joshi *et al*., 2017) have been developed to overcome the root phenotyping bottleneck and have enabled genomic discoveries for seminal RSA traits in durum wheat (Cane *et al*., 2014; Maccaferri *et al*., 2016; Alahmad *et al*., 2019), bread wheat (Atkinson *et al*., 2015), and barley (Robinson *et al*., 2016). However, root phenotypes observed in early growth stages are less representative of the phenotype of mature roots in the field and often have weak to moderate associations (Mace *et al*., 2012; Borrell *et al*., 2014; Maccaferri *et al*., 2016; El Hassouni *et al*., 2018; Alahmad *et al*., 2020). Furthermore, while several studies have identified QTL associated with barley seedling RSA traits under controlled conditions (Chloupek *et al*., 2006; Naz *et al*., 2014; Arifuzzaman *et al*., 2016; Robinson *et al*., 2016; Jia *et al*., 2019) the genetic drivers of mature RSA traits under field conditions remain poorly understood.

Uncrewed aerial vehicle (UAV) technologies are transforming crop phenotyping, providing high-resolution, scalable data for canopy traits (Kyratzis *et al*., 2017). For example, UAV-captured Vegetation Indices (VIs) precisely estimated sorghum leaf area and canopy cover (Potgieter *et al*., 2017; Zhao *et al*., 2021). In wheat, remote sensing enabled accurate prediction of water and nitrogen use efficiency and assessment of foliar temperature and stomatal conductance in maize (Zhang *et al*., 2019; Yang *et al*., 2020; Brewer *et al*., 2022). These technologies offer insights into traits that are difficult to quantify visually, especially when coupled with machine learning, uncovering new opportunities for crop improvement. Wang *et al*. (2023) described two main approaches for the prediction of above-ground biomass using remote sensing. The first integrates remote sensing data into crop simulation models, such as APSIM, to improve the accuracy of biomass prediction. (Hammer *et al*., 2010; Yang *et al*., 2021; Sadeh *et al*., 2024). The second focuses on machine learning models such as partial least squares regression (PLSR) and Random Forest (RF), which leverage UAV data to predict traits accurately. PLSR is a multivariate regression method adept at handling a large number of predictor variables, even when they exhibit a high degree of co-linearity (Wold, 1975). Historically, PLSR has been widely used in remote sensing and spectral datasets with high co-linearity (Robles-Zazueta *et al*., 2021). Similarly, RF is an ensemble learning method that builds multiple decision trees to enhance prediction accuracy and effectively handle complex datasets (Breiman, 2001). Both methods have been widely applied to leverage UAV-captured data to predict above-ground biomass with high accuracy (Han *et al*., 2019; Huntington *et al*., 2020; Masjedi *et al*., 2020). These indirect approaches are non-destructive and repeatable, with decreased susceptibility to human error and improved accuracies. Furthermore, their scalability enhances their applicability in breeding programs and large-scale variety selection initiatives and improves the feasibility of exploring the genetic architecture of complex traits due to increased population size (Smith *et al*., 2021). Consequently, extensive research has explored indirect methods to predict biomass accumulation using UAV phenotyping including the use of multispectral (Liu *et al*., 2019), hyperspectral (Yoosefzadeh-Najafabadi *et al*., 2021), and thermal imagery (Pinto and Reynolds, 2015a).

Some of these studies have explored the potential to use UAV-captured VIs or canopy temperature as ‘proxy’ traits for having improved access to soil water through improved RSA and report promising correlations (Lopes and Reynolds, 2010; Pinto *et al*., 2010; Pinto and Reynolds, 2015a; Li *et al*., 2019). This relationship is potentially driven primarily by leaf surface evaporative area, stomatal conductance and root access to moisture in the soil (Ober *et al*., 2021). Attempting to investigate water supply and demand, earlier work demonstrated the potential for using spectral reflectance in the form of an index linked to water-use and reported significant genetic variation for both leaf water potential and underground water availability in drought conditions (Gutierrez *et al*., 2010). Building on this work, the empirical machine learning approach has yet to be applied to predict RSA. The potential to use VIs to predict RSA could offer advantages for more accurate selection or genetic dissection of RSA-specific genetic controls, as predicted trait values may be less confounded with phenology or other canopy traits across a breadth of genetic diversity.

This study explored the potential of UAV-captured VIs to predict RSA traits and above-ground biomass in barley. We trained machine-learning models on high-quality phenotypes for a representative subset to evaluate prediction accuracy for both RSA and canopy traits. To demonstrate the potential to scale-up the approach, the most accurate model was applied to predict the traits in the broader panel, which were then used for genomic analyses to identify haplotypes associated with above- and below-ground traits. This research highlights UAV technology as a scalable solution for field-based root phenotyping, providing a pathway to develop climate-resilient barley varieties and the potential to extend these applications across diverse crops.

## Materials and methods

### Plant material

This study evaluated a panel of 395 diverse Australian barley breeding lines, representing a broad spectrum of genetic diversity from historic and modern breeding lines and Australian commercial cultivars. The panel was genotyped with 12,561 high-quality single nucleotide polymorphisms (SNPs) using the Infinium™ Wheat Barley 40K v1.0 BeadChip (Keeble-Gagnère *et al*., 2021). This population, which captures the genetic diversity of Australian breeding across the last decade, was split into two experimental testing groups. One group contained the entire panel, while the other consisted of 20 genotypes (Core20) broadly representative of the genetic diversity of the wider panel (97% genetic coverage). Notably, the Core20 subset was carefully selected to share a 5-day flowering window, ensuring it had ideal material for root characterisation without any confounding effects of phenology. These two separate experiments were conducted in Gatton, QLD, Australia (-27.544442, 152.357829), an area with deep clay soils with a high water-holding capacity (Powell, 1982). Population structure analysis was conducted using the “SelectionTools” package in R. Genetic distances between individuals were calculated using modified Roger’s Distance (Rogers, 1972). Classical multi-dimensional scaling based on k-means clustering was applied to the data, followed by hierarchical clustering using Ward’s method (”Ward.D2”) to minimise within-cluster variance (Murtagh and Legendre, 2014).

### Field trial experimental design

The study consisted of two adjacent field experiments, both rainfed with starting irrigation of 30 mm and co-located at The University of Queensland Gatton Research Facility, QLD, Australia (- 27.544442, 152.357829). Experiment one, the ‘Yield Trial’ (YT), consisted of 600 yield plots. A randomised incomplete block experimental design was generated using the Optimal Design package V1.0.1 (Harman and Filova, 2019). The Optimal Design was determined by line allocation within a block and across columns and rows based on relatedness using the additive realised relationship matrix calculated in the ASRGenomics R package (Gezan *et al*., 2022). The entire breeding panel was included in the YT with a partial replication of 1.5 replications (Butler *et al*., 2017; Filov and Harman, 2019). The second experiment, the ‘Coring Trial’, contained plots with the same dimensions and seeding characteristics as the YT. This experiment included the Core20 panel and was designed using Optimal Design as a randomised complete block design with six replicates per line and allocation to row and column position optimised based on the additive realised relationship between individuals.

### Above -and below-ground phenotypes for the Core20

The subset of 20 genotypes was phenotyped for ground-based measurements of above-ground biomass and below-ground root distribution in the field. Above-ground biomass was measured at three timepoints corresponding to key growth stages (GS) depending on the genotype: early tillering (between GS21 and GS25), stem elongation (GS30-GS39), and flowering time (GS50-GS59; Fig. 1). Plants within a 0.25 m^2^ quadrat were hand-harvested at ground level using a sickle, dried at 65°C for seven days, and above-ground dry biomass (SDB) recorded.

**Fig. 1.**
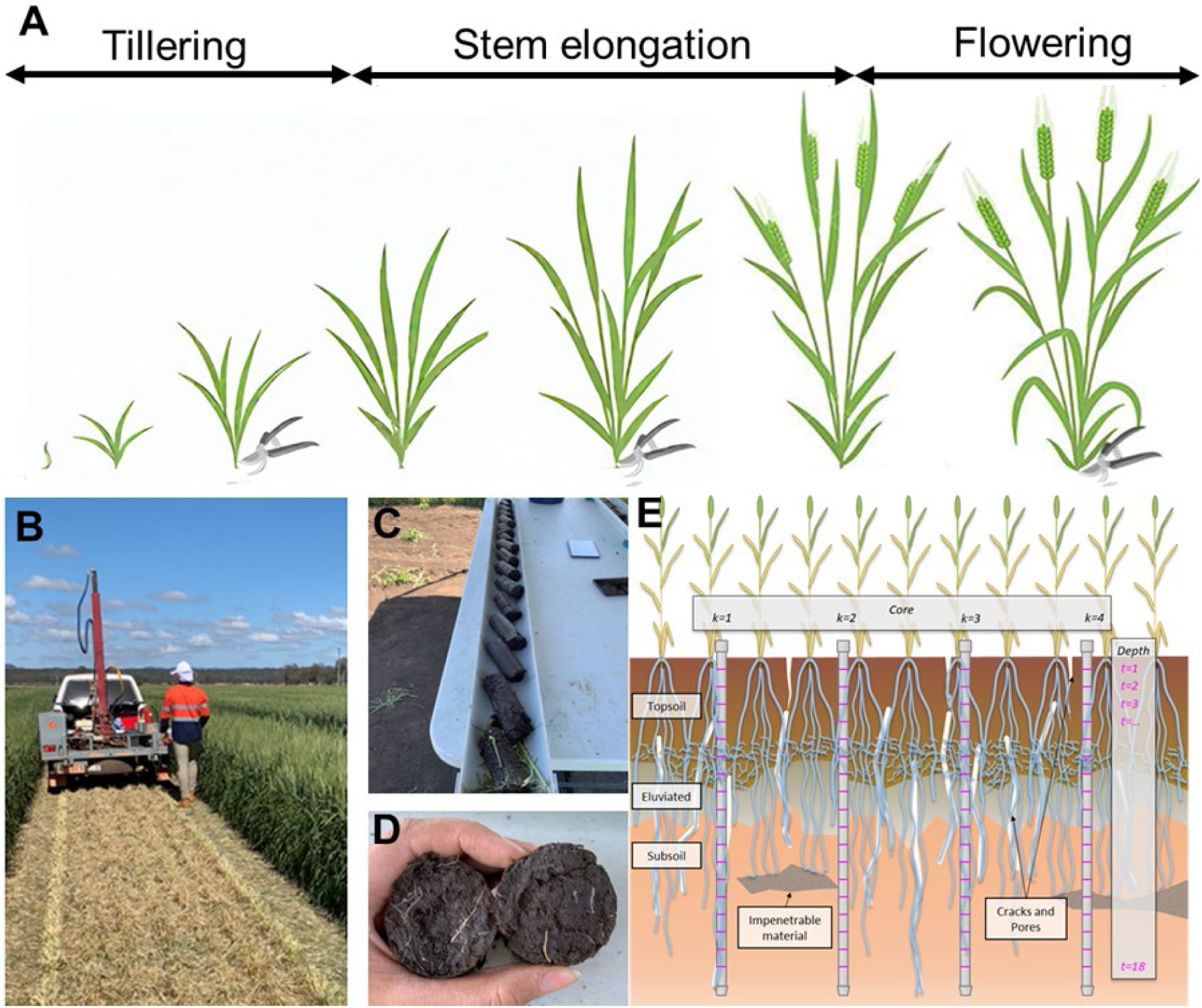
Overview of the high-quality ground-based measurements for above-ground dry biomass and below-ground root count in the Coring Trial. A) Biomass cuts at the three key growth stages: tillering, stem elongation, and flowering time. B) Mulched plots at flowering time in preparation for soil coring using a rig mounted at the back of a trailer. C) Breaking of soil cores into 10 cm intervals. D) Visual counting of roots on each surface of the break. E) Illustration of the position of the four cores sampled within each plot showing the inter- and intra-rows sampling of the core break method as Wasson et al. (2017) described.

The below-ground root phenotyping (root count) was performed at flowering time (GS50-GS59) using the previously reported ‘core-break’ method (Wasson *et al*., 2014). To accommodate for this method, the entire canopy was mulched immediately before soil coring began. Then, a total of 480 soil cores were extracted from the Coring Trial. Four cores were taken per plot, two each from the inter- and intra-rows of plants, avoiding the bordering plants to minimise edge effects. Once the cores were extracted from the ground, they were unloaded and broken into 10 cm intervals (from 0-180 cm), and a visual count of the number of roots visible on each surface of the break was recorded. A subsample of 10 % of the soil cores of two key commercial cultivars (RGT Planet and Maximus CL) was washed to investigate the association between root count and root dry biomass. Root dry biomass (RDB) was recorded after drying at 65°C for seven days.

### UAV phenotyping

To investigate the potential of using UAV-captured VIs as a proxy to explore variation for canopy biomass and root systems indirectly, we performed weekly UAV flights from sowing to flowering for the Coring Trial, while the YT experiment was phenotyped with 17 flights from sowing to physiological maturity. Flight route, image capture, image processing and VIs extraction were performed as described by Smith *et al*. (2024). Briefly, all flights were performed using a Matrice 300 RTK-DJI drone fitted with a ‘MicaSense Altum’ multispectral and thermal high-resolution camera.

Flight altitude was set at 20m with 80% side and front image overlap to ensure sufficient common tie-points during orthomosaic generation. These flight parameters resulted in a ground sample area of 0.86 cm^2^ and 13.5 cm^2^ per pixel for multispectral and thermal, respectively. Flights occurred on clear, still days between 10 am and noon, aligning with the ‘When2Fly’ application to minimise glare and enhance image resolution. (Jafarbiglu and Pourreza, 2023). Further, ten ground control points (GCPs) were placed in each experiment and their accurate GPS coordinates were recorded using Propeller Aeropoints (https://www.propelleraero.com/aeropoints/). The raw images were processed and stitched, using the GCPs to optimise image alignment in Agisoft Metashape software (Agisoft LLC, St. Petersburg, Russia). A single geo-referenced orthomosaic TIF image was generated per flight consisting of six channels, including blue (476 ± 32 nm), green (560 ± 27 nm), red (668 ± 14 nm), red edge (717 ± 12 nm), near-infrared (842 ± 57 nm), and long wave thermal infrared (LWIR) (11 ± 6 µm). Shapefiles containing single plot polygons were created in ArcMap V10.8 to extract plot-level VIs.

Masked and un-masked VIs were calculated for each plot; Masked VIs were obtained by calculating Optimised Soil Adjusted Vegetation Index (OSAVI) and applying Otsu thresholding to the OSAVI-captured pixel values. This method enables the segmentation of green vegetation from soil pixels (Otsu, 1979), and the subsequent calculation of the mean VI value in vegetation-only sections of the plot. In addition, un-masked VIs were also calculated by taking the mean of VI values across the entire plot. All raster calculations and zonal statistics were performed using a Python-based in-house package by researchers from the University of Queensland called XtractoR (Das *et al*., 2022). This tool efficiently extracts a diverse range of VIs and canopy temperature (CT) traits from orthomosaics, with the flexibility to apply soil background masking depending on the target trait. For this study, a total of 60 spectral VIs (Table S1) were calculated as described by Smith *et al*. (2024) in addition to three temperature-related indices.

### Data analysis

Phenotype data, including above-ground dry biomass at each harvest and plot level root counts, were analysed using ASReml-R Version 4.2 (Butler *et al*., 2017) in R version 4.3.0 (R Core Team., 2023). A linear mixed model (LMM) framework was employed to partition various components and account for spatial variation in the field. The model was fitted with genotype as a fixed effect, while replicates were fitted as random effects. Best Linear Unbiased Estimates (BLUEs) for adjusted genotype mean values were calculated using the following Equation 1:

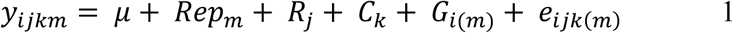

where *y_ijkm_* denotes the plot observation for genotype *i* in replicate*m*, row *j* and column *k*. Fixed effects include the overall mean μ and genotype *G*_*i(m)*_, while random effects accounted for replicate *Rep_m_*, row *R_j_*, and column *C_k_*. The residuals *e*_*ijk*(_*_m_*) were spatially correlated, following *N*(0, *AR*1 ⊗ *AR*1σ^2^), while the variance components for rows and columns were modelled as 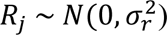 and 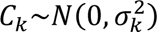, respectively. The adjusted BLUEs were used for subsequent analyses and model training. The broad-sense heritability (H^2^) for root and canopy traits was also calculated.

A semi-two stage modelling approach was performed to gain insights into variation for the overall root system size and distribution at different depths to model root count over depth on a plot level (Pérez-Valencia *et al*., 2022). Packages used as part of this modelling approach included the Spatial Analysis of Field Trials with Splines (SpATS) package (Rodriguez-Alvarez et al., 2018) in combination with the statgenHTP package (Millet *et al*., 2021) in R. Plot level root count BLUEs were used for modelling longitudinal trends using a flexible hierarchical P-spline. This modelling approach allows the calculation of five root proxy traits; (1) area under the root count curve, which is a proxy for overall RSA system size, and (2-5) the area under the curve (auc) at specific depths, namely 0-20cm, 20-60cm, 60-100cm and 100-180cm, as proxies for root distribution in different soil strata (Fig. 3A) Those five root proxies were used in the subsequent machine learning model training and spatial analysis.

Pearson’s correlation (*r*) was calculated using the BLUEs values to assess the relationship between the recorded RDB and root count across different depths of the soil core.

### Training datasets

Three data sets were used to train predictive models: i) high-quality ground-based measurements for above-ground biomass from the three key growth stages (i.e. early tillering, stem elongation and flowering time), ii) precise below-ground root counts using the ‘core break’ method, and iii) UAV-calculated VIs from both field experiments. Through these datasets, we explored the possibility of using UAV-captured VIs to predict above-ground dry biomass and the auc for root number at specific depths, as detailed below.

### Biomass prediction models

Raw-SDB values measured at the three timepoints were assembled alongside the corresponding VIs measured from UAV images captured simultaneously. Subsequently, the data frame (with 165 observations of SDB and 65 predictor VI variables) was split to train and test sets using an 80 / 20 % split. Then, sampling was stratified based on the timing of SDB measurements to ensure even sampling across the three time points. This study used two commonly used machine learning approaches to predict SDB, including PLSR and RF using the Caret package in R (Kuhn, 2008). The first model, PLSR baseline, containing all 63 variables (PLSR_allvars) was used to identify key variables associated with SDB. This model used leave one out cross-validation and a tuning grid of 1:63 for the parameter ‘ncomp’, which defines the number of components in the model. The optimal number of components was chosen based on the minimum Root Mean Square Error (RMSE) achieved across the cross-validation. Subsequently, the VarImp function from the Caret package was used to rank all input variables in PLSR_allvars (Table S1), achieved by taking the weighted sums of variable coefficients across all relevant components (Kuhn, 2008). Starting from the top-ranked variable, we calculated Pearson’s correlation (*r*) between each variable and the next-best-ranked variable. If the correlation was less than a threshold of *r* = 0.8 that variable was also added to the list of final variables until seven predictor variables were selected. The number seven was chosen based on the ‘elbow method’, which iteratively increases the number of variables in a simple multiple linear regression model until a point of inflection in the model variance, which signifies the point of diminishing returns in performance related to the inclusion of any additional variable (Zhang, 2022).

Subsequently, another PLSR model (PLSR_topvars) was created using the seven selected variables to compare model prediction accuracy before and after the variable selection process. In addition to PLSR, we also trained two separate models using RF to estimate SDB. A baseline RF model (Ranger_allvars) was first trained using the ‘Ranger’ package in R, using all 63 variables. Hyperparameter tuning involved iterating through the number of randomly selected features in the model; ‘Split.Rule’ was set to variance, the min.node size was set to 5 and the number of trees was set to 150. A refined RF (Ranger_topvars) was also trained using the top variables selected using the PLSR_allvars. The optimal number of components was chosen based on the maximum coefficient of determination (*R²*) and minimum Root Mean Square Error (RMSE), and Relative Root Mean Square Error (rRMSE) achieved across the cross-validation as detailed in Equations 2-4 below:

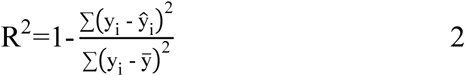

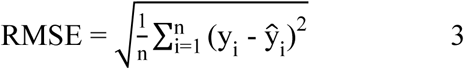

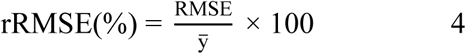

where y_i_ is the observed value for the i^th^ observation, 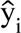 is the predicted value for the i^th^ observation, n is the total number of observations, and 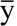 is the mean value of all observations.

To evaluate the model performance, coefficient of determination (R²), Root Mean Squared Error (RMSE), and relative RMSE (rRMSE) were used. Both training and test datasets were utilised to evaluate model performance across different growth stages. Further visualisation, including scatter plots and density curves, were generated to compare observed versus predicted biomass values. Spatial analysis using the R package SpATS was conducted to assess the spatial variability of predicted biomass values and to estimate variance components and heritability of biomass cuts across the entire YT.

### RSA prediction model

RSA traits, including the total auc of the root count growth curve and auc at specific depths (i.e. 0-20cm, 20-60cm, 60-100cm and 100-180cm), were also estimated using the four previous modelling approaches. While the above-ground biomass models utilised VIs calculated at the time of the biomass cuts, VIs measured across the entire season were used to estimate RSA traits. This resulted in a dataset with 550 observations of RSA traits (5 traits x 110 plots), and 1171 predictor variables. The PLSR model using top predictor variables (PLSR_topvar) was chosen to predict RSA traits due to its superior ability to predict RSA traits precisely. Subsequently, the trained PLSR and RF models were applied to predict RSA traits for each root section in each experimental plot for the larger set of individuals in the YT.

The predicted root values were subject to spatial analysis using the SpATS package in R. Variance components were computed to identify the proportion of variation explained by genetic factors, which enabled the calculation of broad sense heritability for all RSA traits.

### Genome-wide haplotype approach for predicted canopy and RSA traits

A total of 12,561 SNP markers were subjected to quality control, and full marker profiles were provided following imputation using Beagle version 5.4 software (Ayres *et al*., 2012). This included the removal of markers with minor allele frequency (≤0.05) and those with greater than 10% heterozygosity, resulting in 6,753 high-quality polymorphic SNPs. (Gower, 1966; Murtagh and Legendre, 2014)

A local genomic estimated breeding value (LGEBV) haplotype mapping-based approach was carried out (Voss-Fels *et al*., 2019). Phenotypic data used in this approach was BLUEs calculated for the predicted five RSA traits and the three canopy SDB using PLSR_topvar model applied to the entire population of 395 breeding lines. Initially, linkage disequilibrium (LD) blocks were assigned using the ‘Selection Tools’ package, based on patterns of LD present in the marker data. A total of 2,593 LD blocks were classified by applying an LD threshold of 0.6, a marker tolerance of 3, and an average of 2.6 markers per block. For each haplotype, LGEBV was calculated by first determining the individual SNP effects using a ridge-regression best linear unbiased prediction model (rrBLUP). Then, the SNP effects within each block were summed and the haplotype effects were calculated within the block. The variance for haplotype effects within each LD block was calculated, and the top 0.5% of blocks with the highest variance were deemed significant. To facilitate the visualisation of the significant blocks associated with SDB and RSA traits, we used CMplot version 4.5.1 (Lilin-Yin, 2023). The physical start and end positions of the top 0.5% high-variance haploblocks were identified using Morex V1 as a reference and compared against known QTLs and genes regulating RSA traits, with a linkage map generated in MapChart (Voorrips, 2002) to assess overlaps with developmental QTLs for above- and below-ground traits (Fig. S4).

## Results

### Population structure

Clustering analyses identified three distinct hierarchical clusters. Cluster 1 represents an Australian germplasm pool 1 originating from ancestral south-eastern Australian breeding programs, and includes modern varieties such as Spartacus CL, Maximus CL, and Compass. Cluster 2 consists of European-derived barley lines and varieties such as RGT Planet and Oxford. Cluster 3 encompasses a second Australian germplasm including the varieties Buff, Combat, and Scope CL, which are derived from either the ancestral Western Australian and Victorian barley breeding germplasm pool or the more recent InterGrain breeding outcomes. A PCA revealed that 62.01% of the total variance was captured across 10 Principal Components (PCs), with the first two PCs accounting for 12.4% and 11.6% of the variance, respectively (Fig. 2).

**Fig. 2.**
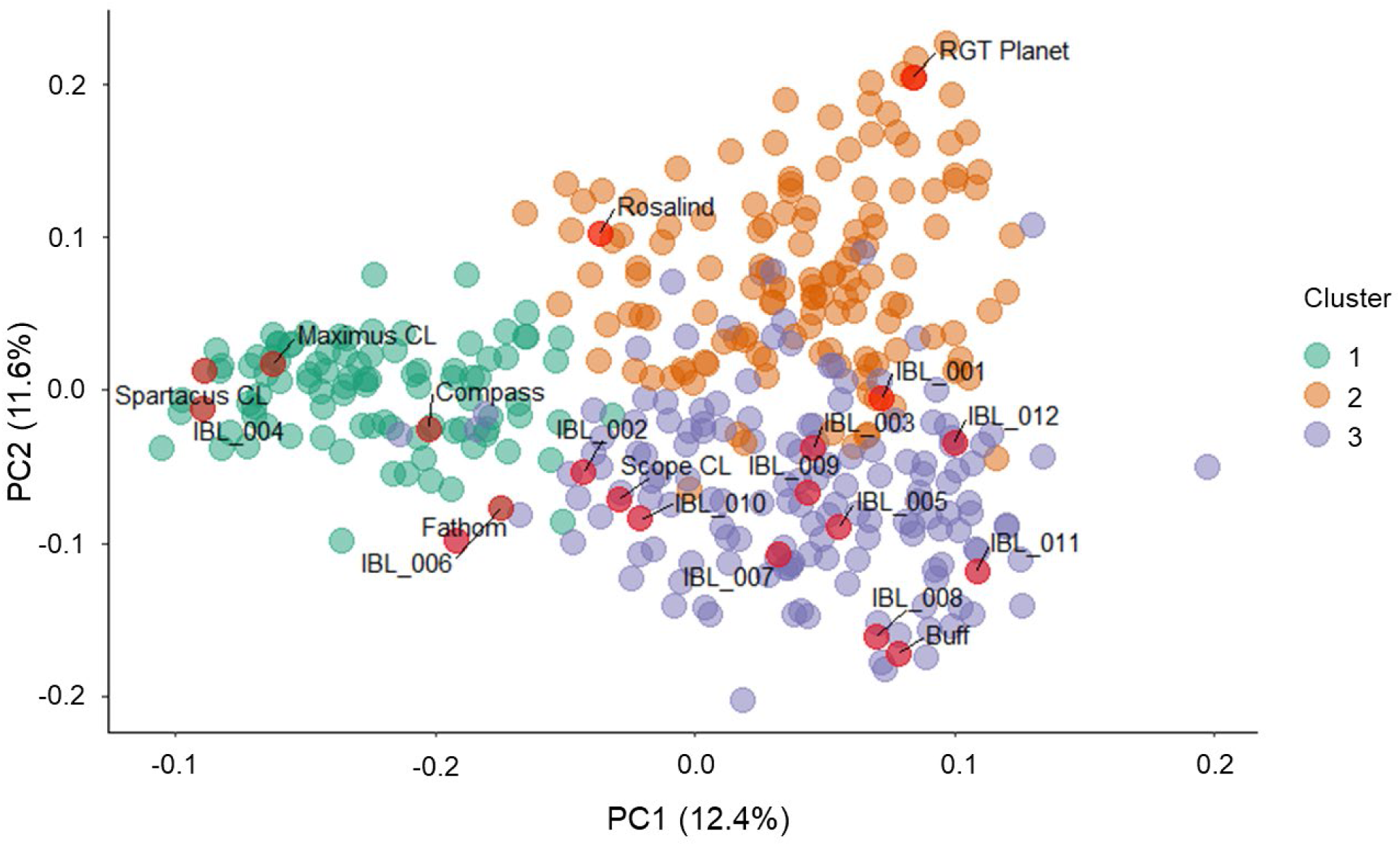
Population structure of the Australian barley breeding panel, calculated from 6,753 polymorphic SNP markers for 395 lines. Three main clusters were identified: Cluster 1 consists of Australian germplasm pool 1, including germplasm from the VIC and SA breeding programs (green). Cluster 2 represents European-derived barley breeding lines and varieties (orange). Cluster 3 comprises Australian germplasm pool 2, which includes lines that are descendants of acid-tolerant material, generally originating from the historic WA and QLD breeding programs (violet). Genetic distance was calculated, with the variance explained by principal components being PC1 = 12.4% and PC2 = 11.4%. The subset of 20 lines used in the Coring Trial are presented in red and represent 97% of the Australian barley breeding panel.

### Variation for canopy and root traits across 20 representative lines

A subset of 20 lines was subjected to ground-based phenotyping for above-ground biomass and below-ground root distribution, providing training datasets for machine learning models to predict these traits in the larger set of 395 breeding lines (Fig. 3). SDB measurements collected for the training set at early tillering, stem elongation, and flowering time (Fig. 1), displayed a consistent SDB increase over time, with a substantial range observed across the three timepoints. For instance, the mean adjusted SDB value for early tillering was 37 g/m^2,^ with a range spanning from 24.5 g/m^2^ to 49.0 g/m^2^. For SDB during stem elongation, the adjusted mean value was 129.9 g/m^2^ with a range spanning from 74.9 g/m^2^ to 215.6 g/m^2^, and the SDB adjusted mean value at flowering time was 305.6 g/m^2^ with a range of 191.8 g/m^2^ to 425.0 g/m^2^ (Fig. 3B).

**Fig. 3.**
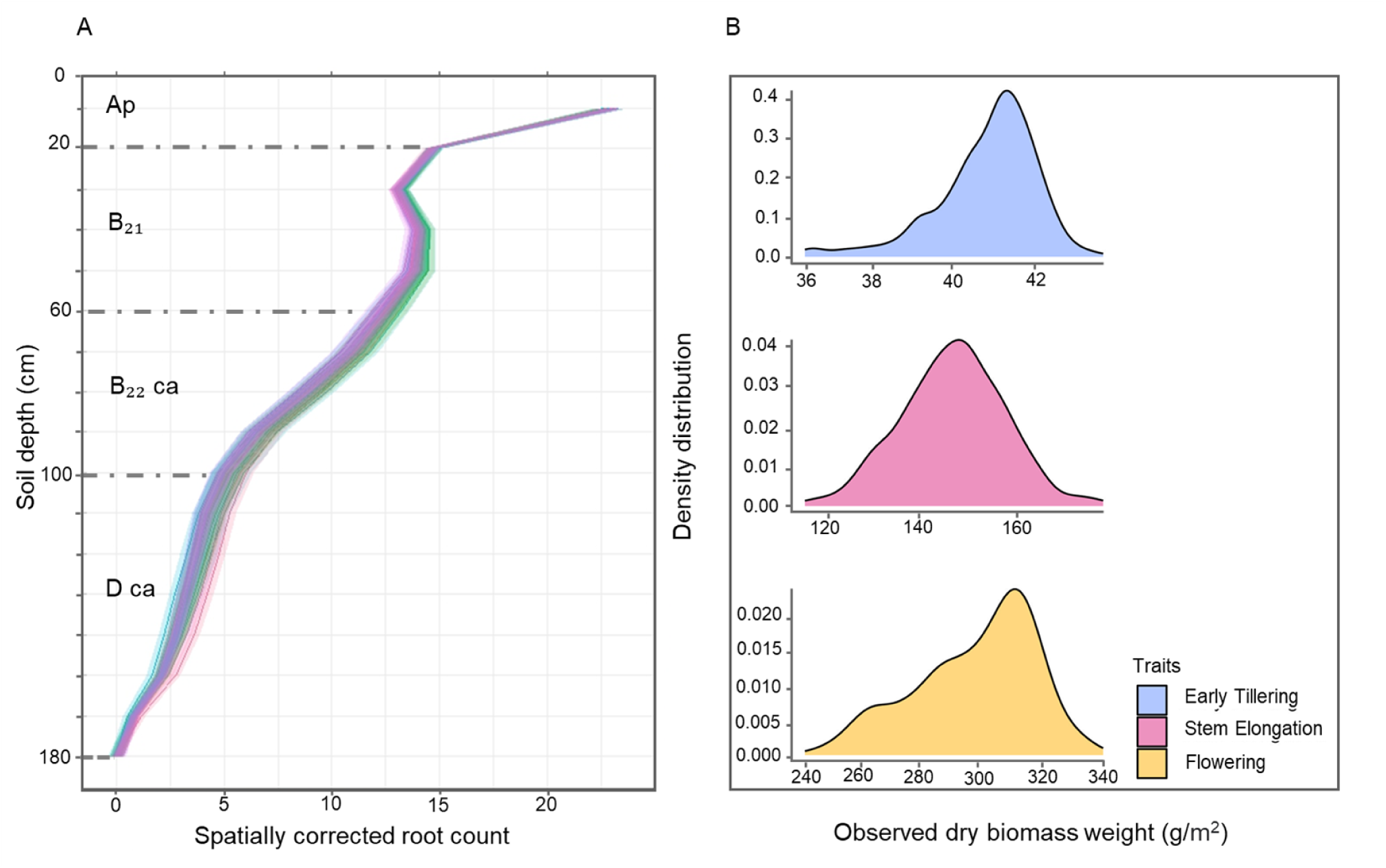
Root distribution profile and density distribution of above-ground biomass for the 20 lines used as a training population for phenotypic predictions. A) Spatially corrected root count for the different soil depths collected at flowering time, fitted by a spline-model; horizons from the soil are presented (Powell, 1982): 0-20cm (Ap horizon, brownish-black clay, pH 6.5-7.8), 20-60cm (B21 horizon, brownish-black heavy clay, pH 8.2-8.8), 60-95cm (B22 ca horizon, brown clay with carbonate, pH 8.2-8.8), and 95-150cm (D ca horizon, brown light clay with carbonate, pH 8.2-8.8). B) Density distributions of above-ground dry biomass (g/m²) at three growth stages: Early Tillering, Stem Elongation, and Flowering.

Plot level root counts showed considerable variability for root distribution across different soil strata, with a progressive decrease in root counts observed from the upper to the deeper layers of the soil (Fig. S1). At 10 cm depth, the mean root count was 23.2, gradually decreasing to 0.4 at 170 cm depth. Genotypic variation in root distribution was observed within each root section, while greater variability was observed within the upper layer of the soil. For instance, at 10 cm depth, the range of root counts varied from 10.5 to 38.75. Five root proxy traits were calculated - the total auc of the root count curve and the auc of root count curve at various soil depths (Fig. 3A), and significant genotypic variation was observed. For instance, for the area under the curve at the 0-20 cm depth interval, values ranged from 17.7 to 19.9, with an average of 18.2. In the deeper sections of the soil the auc increased significantly, with the highest average auc observed in the 20-60 cm depth (41.1). However, the auc for 100-180 cm soil depth experienced a dramatic decrease in root growth and distribution compared to the other RSA traits. The overall auc value, representing the cumulative root activity across all depth intervals, ranged from 90.6 to 192.9, with an average of 128.1 (Fig. 3A).

### Relationship between root count and root biomass in barley genotypes

Overall, higher root number and more RDB were observed in the RGT Planet in comparison to Maximus CL as displayed in the marginal density distributions (Fig. 4). Regression analysis indicated a significant positive association between root count and RDB (*r* = 0.63, *p* < 0.001***), consistent with the expected correlation between root count and biomass through their shared relationship with root length. These results underscore the potential of using root count as a proxy for assessing root biomass.

**Fig. 4.**
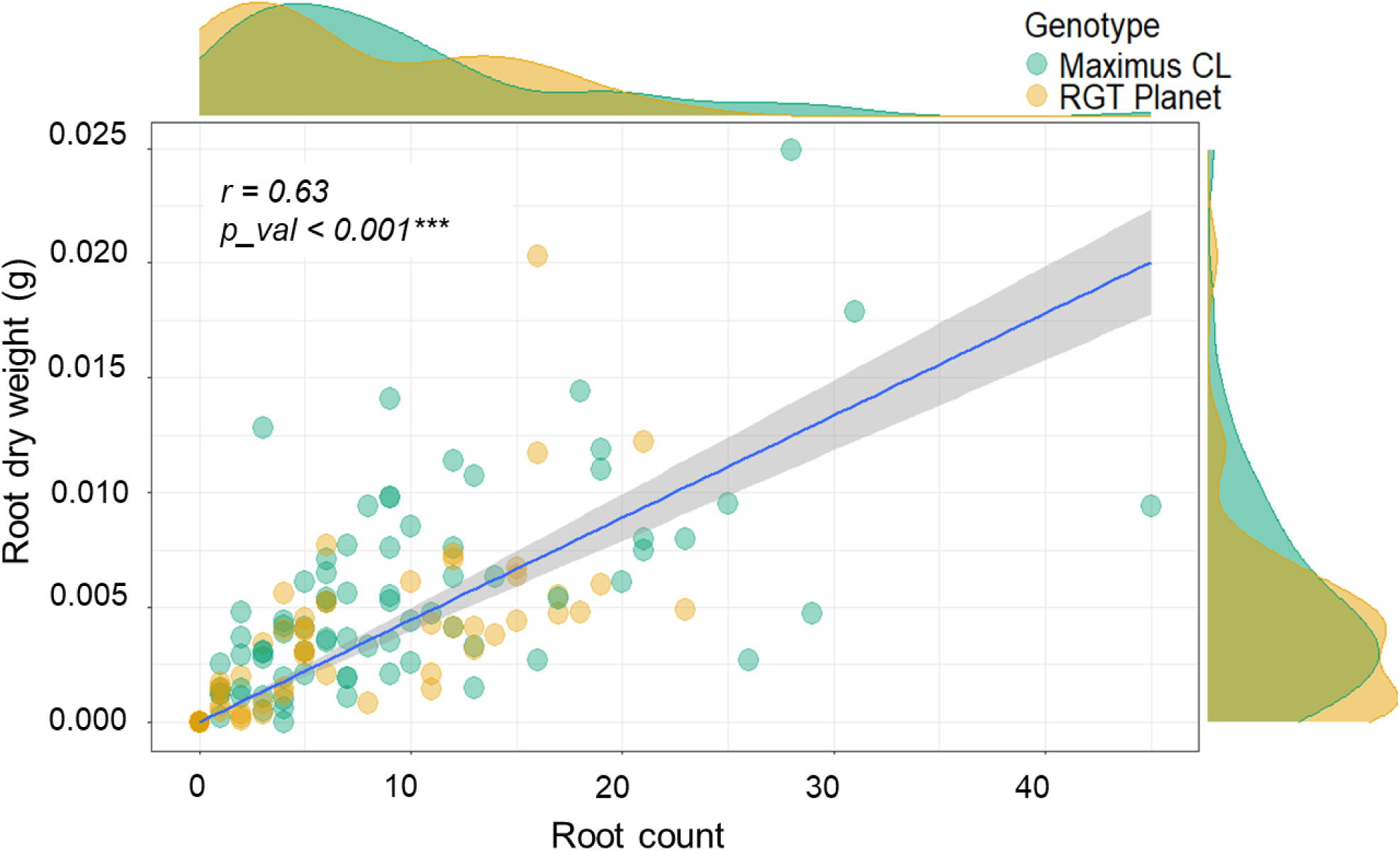
Positive association between root count and root dry biomass (g) in each section of the soil core for Maximus CL and RGT Planet. Each point represents a soil core section (i.e. at a specific depth), and the colour represents the genotype. The Pearson’s correlation (*r*) value is displayed, with significance levels indicated by (*). Marginal density distributions are presented on the axes to reveal the distribution of root count and root dry biomass (g).

### UAV traits enable the prediction of shoot biomass at key growth stages

UAV-captured VIs accurately predicted SDB across three growth stages for 20 genotypes (Fig. 5), enabling the use of SDB data to train PLSR and RF models with 63 high-correlation (≥0.8) variables to improve biomass prediction accuracy. Key variables were identified through PLSR reduced redundancy and maximised predictive power, and a subsequent PCA revealed that 90.9% of biomass variation over time was explained by PC1 and PC2 (Fig. S2). Following this, refined PLSR and RF models were used, utilising the top seven variables identified using the PLSR_allvars. Notably, variables such as Modified Soil-Adjusted Vegetation Index (MSAVI) and osavi-masked mean temperature were identified as key VIs for biomass prediction and had a significant negative association with biomass at flowering time (*r* = -0.97, *r* = - 0.72), respectively. A strong correlation between observed SDB and predicted SDB at the three growth stages was found in both training and testing datasets using PLSR and RF models. However, the final model used for biomass prediction of the entire population (*n* = 395) was the PLSR model with seven variables, which demonstrated improved accuracy and consistent performance (Fig. S2). For instance, in the training set, the model exhibited an RMSE of 22.6g/m^2^, a rRMSE of 14% and a R^2^ of 0.98; for the independent testing set, an RMSE of 21.9 g/m², rRMSE of 15.2%, and R^2^= of 0.98 was observed (Fig. 5A). Predicted SDB exhibited a significant variation across the three growth stages. For instance, the mean predicted SDB was 40.79 g/m² during Early Tillering, 147.31 g/m² at Stem Elongation, and 298.05 g/m² at Flowering (Fig. 5B).

**Fig. 5.**
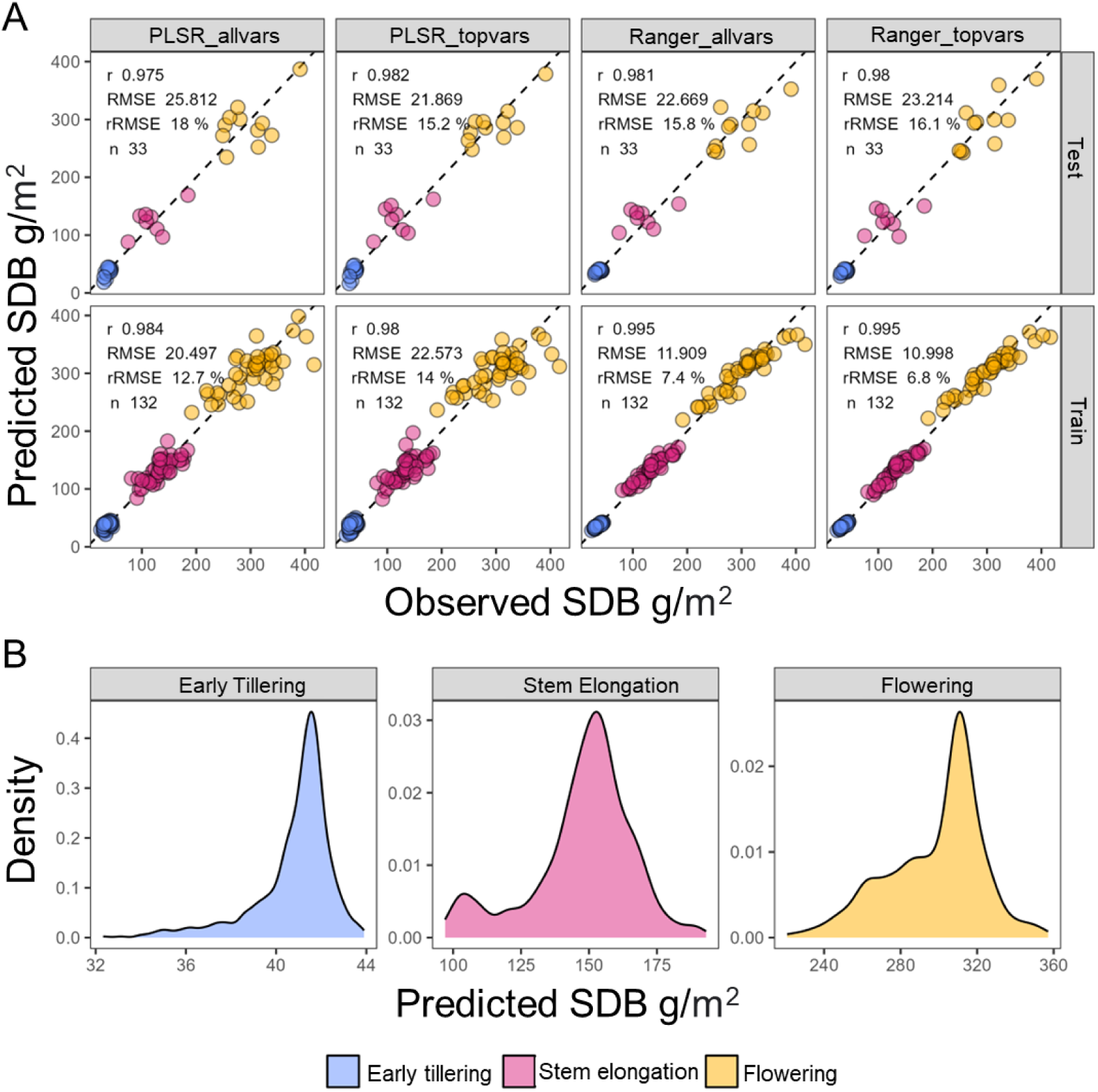
Predicted above-ground dry biomass with four modelling approaches. Partial Least Square Regression using all vegetation indexes (PLSR_allvar), PLSR using the most important variables (PLSR_topvars), Random Forest using all variables (Ranger_allvars), and Random Forest using the most important variables (Ranger_topvars). A) Observed versus predicted above-ground dry biomass using UAV vegetation indexes calculated at three-time points. The horizontal X-axis represents the predicted SDB for the set of 20 genotypes obtained from the model, and the vertical Y-axis represents the SDB measured manually at ground level. Correlation (*r*), residual mean square error (RMSE), ratio residual mean square error (rRMSE) and the number of individuals (*n*) are also displayed for each model. B) Density distribution displaying the variation of the predicted biomass for the panel of 395 Australian barley breeding lines using VIs at early tillering, stem elongation, and flowering stages and PLSR_topvars model.

### Vegetation indices captured by UAV enable the prediction of root traits in the field

UAV-captured VIs, initially utilised to predict SDB at three growth stages (early tillering, stem elongation, and flowering), were also employed to predict RSA traits. These traits included the auc of the root count growth curve and auc at specific depths corresponding to different soil strata, as detailed in Fig. 3. The PLSR model was chosen to predict RSA traits for 395 genotypes. PCA highlighted VIs that influence RSA trait prediction (Fig. S3). Using the seven selected variables by PLSR model, the PC1 explained (78.6% - 88.6%) variation and PC2 explained (3.8% - 14.3%) variation for all RSA traits (Fig S3). Interestingly, the mean temperature at 88 days after sowing (‘mean_temperature_88’) was a key predictor of RSA distribution and size. The model’s performance varies for RSA traits across different soil depths. For the training set, the overall auc showed an RMSE of 13.2, an rRMSE of 10.3%, and an R² of 0.719. The 0-20cm depth range showed the lowest error with an RMSE of 0.266, rRMSE of 1.5%, and R² of 0.507. For the 20-60 cm range, the model showed an RMSE of 2.84, rRMSE of 6.9%, and R² of 0.702. The 60-100 cm depth displayed an RMSE of 3.809, rRMSE of 15.9%, and R² of 0.702. Finally, the deepest range of 100-180 cm demonstrated the highest relative error with an RMSE of 5.242 g/m², rRMSE of 33.7%, and R² of 0.668 (Fig. 6A).

**Fig. 6.**
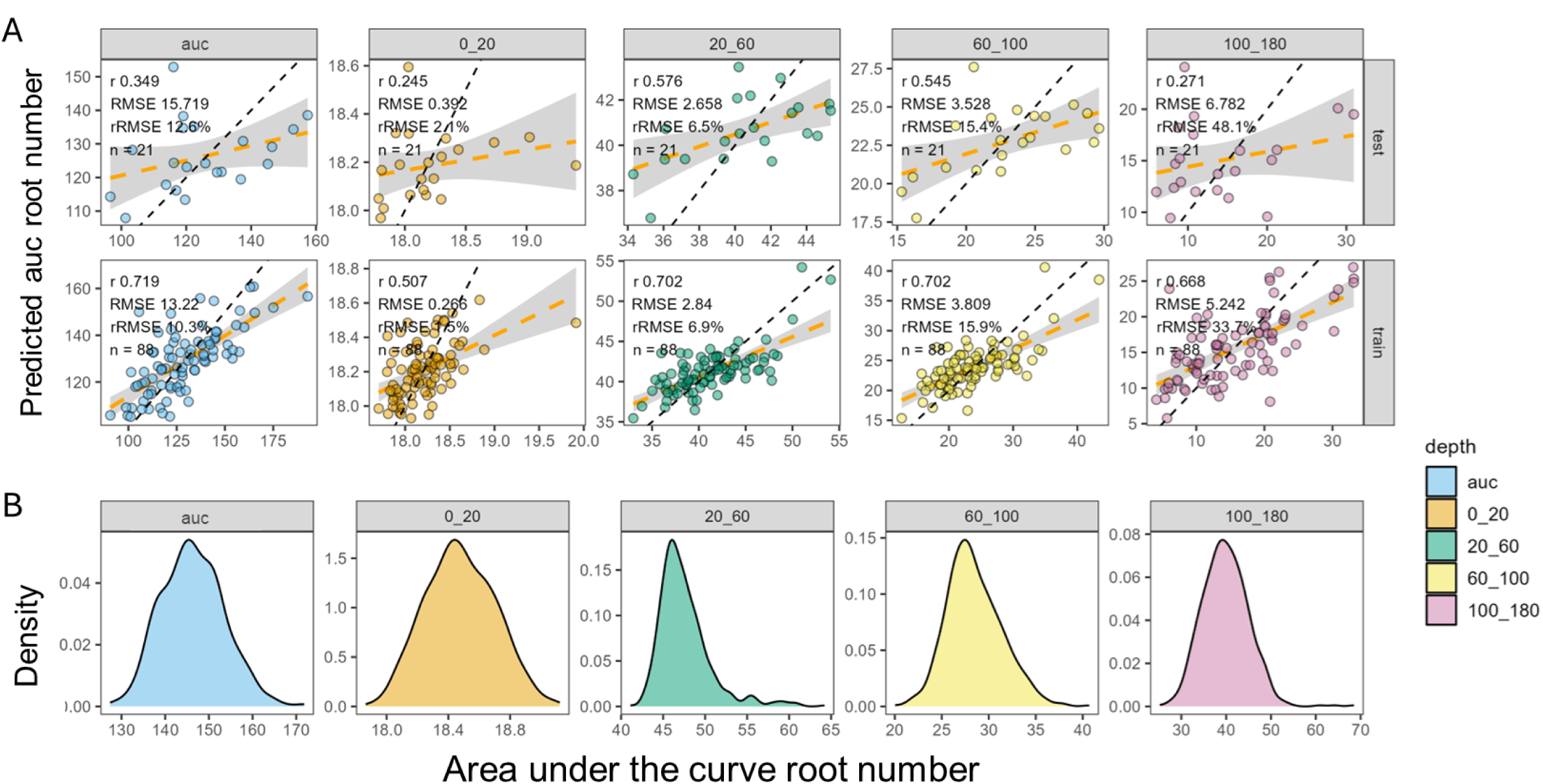
Observed versus predicted overall area under the curve of the root count growth curve (auc) and for specific depth sections for the set of 20 individuals using UAV vegetation indexes. A) The training set (80% of samples) and testing set (20% of samples) were evaluated using the Partial Least Square Regression model with the most important variables (PLSR_topvars). Correlation (*r*), residual mean square error (RMSE), ratio residual mean square error (rRMSE), and the number of individuals (*n*) are displayed for each RSA trait (auc, 0-20, 20-60, 60-100,100-180). B) Density distribution showing the variation of the predicted biomass for the panel of 395 Australian barley breeding lines using the PLSR_topvars model, categorised by root depth.

Predicted RSA traits varied significantly across plots, demonstrating a range of values for different RSA traits corresponding to the modelled overall area under the curve and auc at specific depths (Table S2, Fig. 6B). Broad sense heritability (H^2^) of 0.65, 0.78, 0.74, 0.74, and 0.66 were calculated for overall auc, 0-20cm, 20-60cm, 60-100cm, 100-180cm, respectively.

### Identified haploblocks influence barley root traits and canopy growth

The top 12 haplo-blocks for each trait were identified as blocks of high importance, with 19 of these influencing RSA traits only, and nine exclusive to SDB traits (Table S3). Interestingly, 15 haplo-blocks were identified as common across RSA and SDB traits. Furthermore, five haplo-blocks influenced shallow root depth (0-20cm), while three blocks were linked to the overall RSA size (auc). Regarding canopy development, five blocks exhibited strong associations with SDB at early tillering (SDB29), and two blocks were notably linked to biomass at flowering (SDB81). Interestingly, all 12 haplo-blocks identified for SDB54 were also associated with RSA traits, indicating a strong association between biomass accumulation during the stem elongation growth stage and RSA traits around flowering. Of particular interest, certain haplo-blocks were associated with SDB at the three growth stages early tillering, stem elongation, flowering time and shallow RSA traits (e.g., haplo-blocks b000827 and b000828) on 3H, as well as deep RSA traits (e.g., haplo-block b001434) on 5H. Moreover, haplo-block b001441 was identified as one of the most important blocks controlling all RSA traits, including overall RSA size and depth-specific blocks (auc, 20-60cm, 60-100cm, 100-180cm) on 5H. The significant blocks are presented in (Fig. 7).

**Fig. 7.**
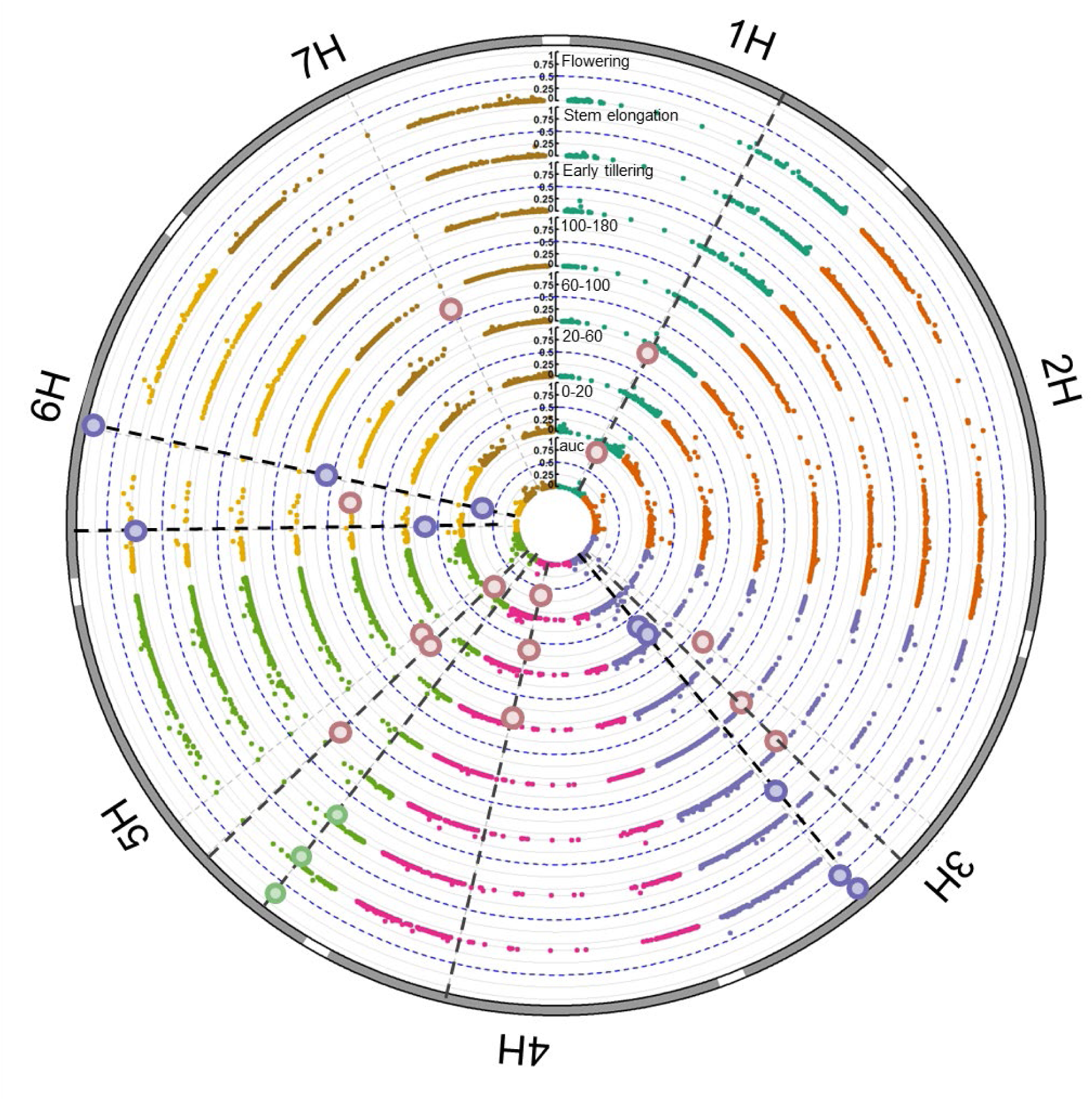
Circos Manhattan plot of haploblocks associated with above-ground biomass and RSA traits in barley. Concentric rings represent the area under the curve (auc) for root count (overall and at various soil depths) and above-ground dry biomass at key growth stages. Dotted lines indicate shared genetic regions for root and shoot traits (blue), canopy traits only (green), and RSA traits only (brown). Barley chromosomes (1H-7H) are labelled on the outer circumference. Only 0.5% of haplo-blocks are considered significant and are presented.

The top 0.5% of haploblocks identified in this study were also aligned with previously reported QTLs or developmental genes in the literature (Fig. S4: Table S4). Among the identified blocks for predicted RSA traits, 14 were novel and four aligned with previously reported genes and QTLs (Diab *et al*., 2004; Chen *et al*., 2010; Kirschner *et al*., 2021; Fusi *et al*., 2022; Siddiqui *et al*., 2024). Additionally, ten haplo-blocks were unique to SDB, seven being novel. Some of the blocks overlapped with already reported QTLs; haploblock b000077, which was strongly associated with SDB29 at early tillering, aligned with the relative water content QTL (*QRWC*) on chromosome 1H, as reported by Chen *et al*. (2010) in a study of a double haploid population under water stress and irrigated conditions. Of particular interest is haploblock b002045 on chromosome 6H, which was strongly associated with predicted RSA traits (60-100cm), aligned with the *EGT1* gene (Enhanced Gravitropism 1) by Fusi *et al*. (2022). Similarly, haploblock b001441 on chromosome 5H, also associated with several predicted RSA traits in our analysis, aligned with the *EGT2* (Enhanced Gravitropism 2) gene which controls gravitropic setpoint angle in barley RSA reported by Kirschner *et al*. (2021).

Moreover, RSA block b002386 on chromosome 7H, linked to the area under the curve measurements for shallow RSA at 0-20cm and deep RSA at 60-100cm and 100-180cm, was associated with the water-soluble carbohydrate QTL at 100% relative water content (WSC100 40) identified by Diab *et al*. (2004). Importantly, haploblock b000387, associated with SDB29, also aligned with three RSA crossing QTLs (*NRC*) on chromosome 2H, as reported by Siddiqui (2024) in a study investigating the RSA stress plasticity index within a spring barley diversity panel. Furthermore, the first leaf length trait (*QL1L.7H*) in a double haploid population studying seedling drought resistance in wild barley (*Hordeum spontaneum*) was aligned with the shallow RSA haploblock b002528 on chromosome 7H.

## Discussion

This study goes beyond traditional UAV phenotyping applications on above-ground traits by using a large number of UAV-captured VIs, in combination with machine learning algorithms, to directly predict below-ground RSA traits. For instance, earlier studies have demonstrated UAV phenotyping capacity in predicting above-ground biomass and water use efficiency in wheat and sorghum using a single VI, such as MSAVI or leaf area index, as indirect proxies for access to soil water (Lopes and Reynolds, 2010; Pinto and Reynolds, 2015b; Potgieter *et al*., 2017). Here, we used 60 spectral VIs and three canopy temperature-related traits across different growth stages to directly predict RSA. By integrating machine learning with ground-truth RSA data, we have developed a scalable solution to phenotype RSA and canopy traits simultaneously, which has the potential to advance research and help address the bottleneck of RSA in large populations under field conditions.

Substantial variation was observed for SDB and RSA traits across the studied barley population, providing a critical foundation for understanding the genetic makeup of these traits. This variation was consistent across different growth stages for SDB and for RSA traits during flowering time. SDB ranged from 24.5 g/m² during early tillering to 425.0 g/m² at flowering, highlighting the influence of both developmental progression and genetic variation among individuals on biomass accumulation, as displayed in the density distribution (Fig. 3B). Furthermore, root counts ranged from 10.5 to 38.75 at 10 cm soil depth, with a greater concentration of roots in the upper soil layers and a gradual decrease in root density at greater depths. Similarly, variation for RSA was also reported for two populations of barley inbred lines over two consecutive field seasons, emphasising the role of variation in understanding the genetic architecture of RSA traits as a target for drought adaptation (Siddiqui *et al.,* 2024). This variation also includes resource acquisition. For example, the concentration of roots in the upper soil layers observed in our study aligns with findings linking nutrient availability to root density. It highlights RSA traits’ role in enhancing water uptake and decreased canopy temperature using UAV phenotyping technology in wheat (Lopes and Reynolds, 2010). Furthermore, observed above- and below-ground variation among the Core20 subset (120 experimental plots) is important for training and testing machine learning models to ensure robust model development for accurate trait prediction of the 395 breeding lines. Future research integrating field experiments and modelling (Manschadi *et al*., 2006; Rich *et al*., 2016) could validate the value of specific RSA traits, such as deeper roots for drought resilience or shallow roots for nutrient uptake (Henry *et al*., 2010; Lynch, 2019), strengthening their role in breeding for environmental adaptation.

LGEBV analysis performed on the 395 breeding lines further demonstrated the utility and value of this approach by enabling the identification of haploblocks associated with predicted SDB and RSA traits under field conditions. Notably, key haploblocks associated with RSA traits on chromosomes 5H and 6H were co-located with *EGT1* and *EGT2* genes. These genes play a critical role in regulating barley roots’ gravitropism, influencing root angle and may influence access to deep soil moisture (Fusi *et al*., 2022). The alignment of haploblock b002045 with *EGT1* and b001441 with *EGT2* represents a robust validation of the method’s ability to dissect the genetics underlying RSA traits under field conditions. This finding is novel compared to earlier studies on barley RSA traits, as no previous research has linked these genes to RSA traits in the field. Therefore, this approach delivers a new opportunity for the root research community to explore the genetics of root systems across diverse environments and assess their contributions to grain yield under water-limited and optimal conditions. In addition to *EGT1* and *EGT2*, several novel haploblocks were identified. For example, haploblocks b000827 and b000828 were associated with RSA in shallow soil strata (0–20 cm) and SDB during stem elongation, suggesting these haploblocks may contain loci that have similar role to *PIN* genes in sorghum that are known to control allometric responses between root and shoot development (Borrell *et al*., 2022). These findings highlight the power of the UAV phenotyping approach to identify both known and novel genomic regions that govern RSA traits under field conditions. This has been a challenge in previous studies focused on canopy traits alone.

The results underscore the broader implications for breeding programs by providing high-throughput screening for RSA traits in parallel with canopy-related traits. While traditional root phenotyping methods is labour intensive, destructive and limited to a small number of genotypes, UAV-based phenotyping offers a scalable, non-destructive solution. However, it is crucial to understand the value of RSA traits in diverse environmental contexts to effectively include these traits in breeding program selection strategies. This remains a somewhat unresolved research question for many corps. Simulation studies have successfully reported enhanced yield in wheat when the crops have an improved ability to access soil water during grain filling (Manschadi *et al*., 2006; Christopher *et al*., 2013). The data generated from our study could similarly be used to improve the prediction accuracy of simulation models such as APSIM and evaluate the impact of RSA traits on yield in different scenarios (Hammer et al., 2010). Predicted RSA traits could also enhance genomic selection models by improving prediction accuracy and enabling plant breeders to evaluate RSA contribution to yield and better understand genotype by environment interactions. Furthermore, predicted RSA traits could be integrated into phenomic selection indices alongside canopy traits to assist breeders in developing varieties tailored to specific target environments.

This study lays a novel framework for future field-based root trait prediction. A limitation of this study is a single season of data, but the success of this work highlights the opportunity for the research community to adopt and adapt the approach for their crop of interest across diverse environments and production scenarios. The accuracy of RSA predictions may vary with soil type, seasonal moisture levels, and across different environments and crops, highlighting the need to explore these variables in future studies. Additionally, although this study leveraged high-resolution UAV imagery and advanced machine learning algorithms, improvements in UAV sensor resolution and further refinement of model parameters could enhance predictive accuracy, especially in crops with different canopy structures (Zhao *et al*., 2022).

In conclusion, this study represents a significant step forward in crop phenotyping by integrating UAV-captured VIs with machine learning to evaluate RSA traits for large experimental or breeding panels. Unlike earlier studies that primarily relied on single VIs, such as NDVI or canopy temperature, to indirectly measure or correlate with RSA traits, this approach employed a comprehensive set of VIs obtained through multiple UAV flights, enabling the direct prediction of traits at different soil depths. The high accuracy achieved in predicting SDB and RSA proxy traits highlights the potential of this high-throughput, non-destructive methodology for root-based field phenotyping. Moreover, the detection of haploblocks containing functional root genes, including *EGT1* and *EGT2*, showcases the potential for this method to perform robust RSA phenotyping at a population scale required for haplotype mapping or genome-wide association studies. From a crop breeding perspective, the scalability has the potential to facilitate evaluation of large breeding populations to support selection for desirable RSA traits which could accelerate genetic gain for adaptive trait combinations required for climate resilient varieties.

## Supplementary data (brief for each item)

Table S1: Spectral indices and temperature variables ranked by correlation with dry biomass.

Table S2: Summary of predicted root system architecture (RSA) traits.

Table S3: Haplo-blocks in barley classified by traits, with unique (black) and shared (red) blocks.

Table S4: QTLs and genes in barley aligned with haplo-blocks, including locations and contexts.

## Acknowledgements

The authors thank the technical and breeding staff at InterGrain Pty. Ltd. for providing the germplasm and genotypic data used in this study. The authors thank the technical team at the University of Queensland for their assistance in trial management.

## Author Contributions

SA and LH designed the experiments. SA, DS, CK, SVM, LM, AW and HR conducted experiments, UAV phenotyping, and data processing. SA and DS analysed data, with ZA aligning root and shoot data to prior studies. SB, JG and DM supplied germplasm. SA drafted the paper, with DS, HR, and LH providing feedback. DS, CK, ZA, SVM, LM, KC, SC, ABP, AW, SB, JG, and DM reviewed it. All authors approved the final manuscript.

## Data availability

Data supporting this study will be made available by the corresponding authors upon request and approval from InterGrain Pty. Ltd.

## Supplemental materials

**Fig. S1.**
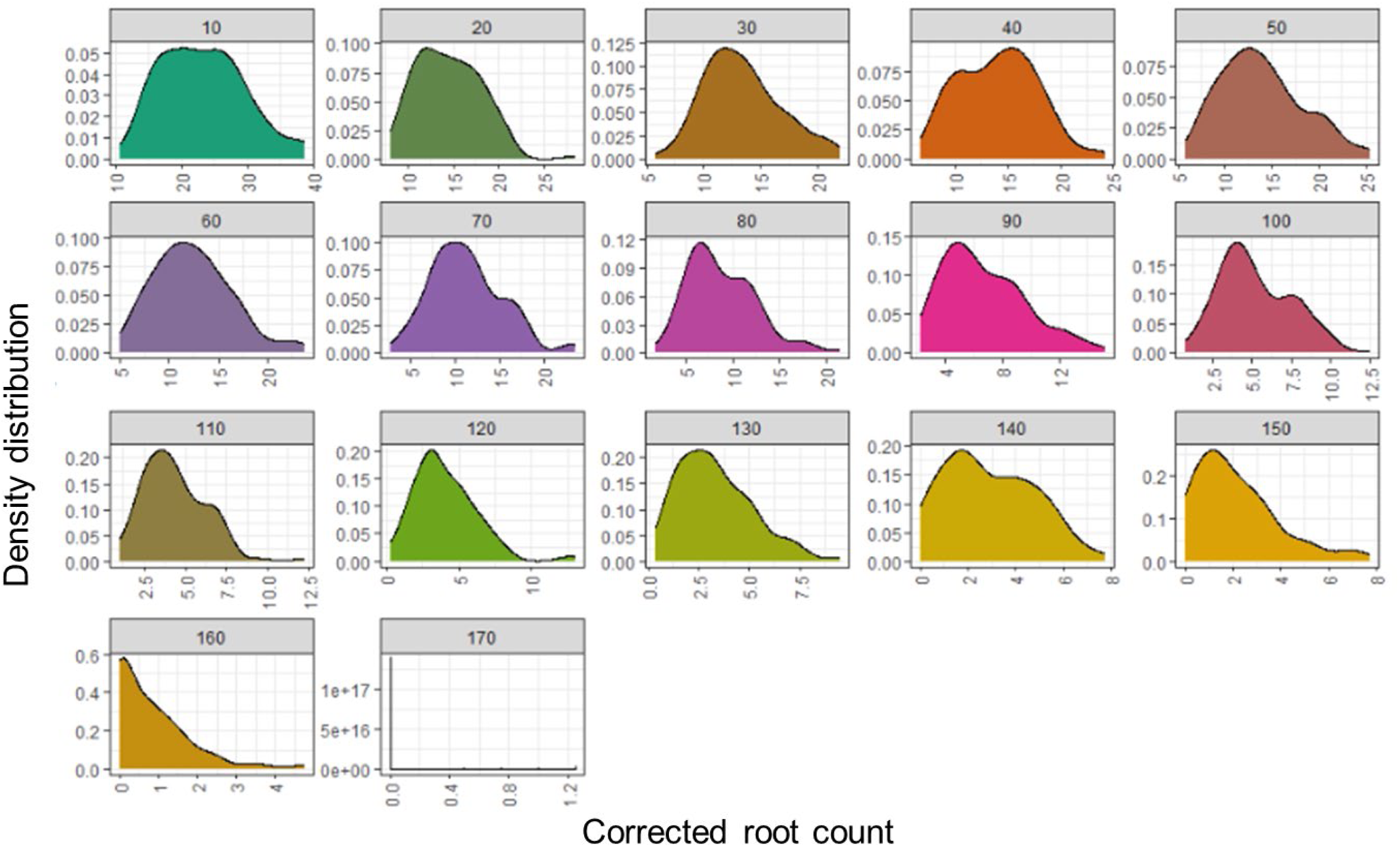
Root number density distribution across soil depth for 20 lines in the Coring Trial at soil depths from 10 cm to 170 cm, in 10 cm intervals. The graph displays adjusted root count density at various soil core sections, showing a decreasing trend in root abundance with depth. Deeper soil layers exhibit a leftward skew in the distribution, reflecting fewer roots at greater depths.

**Fig. S2.**
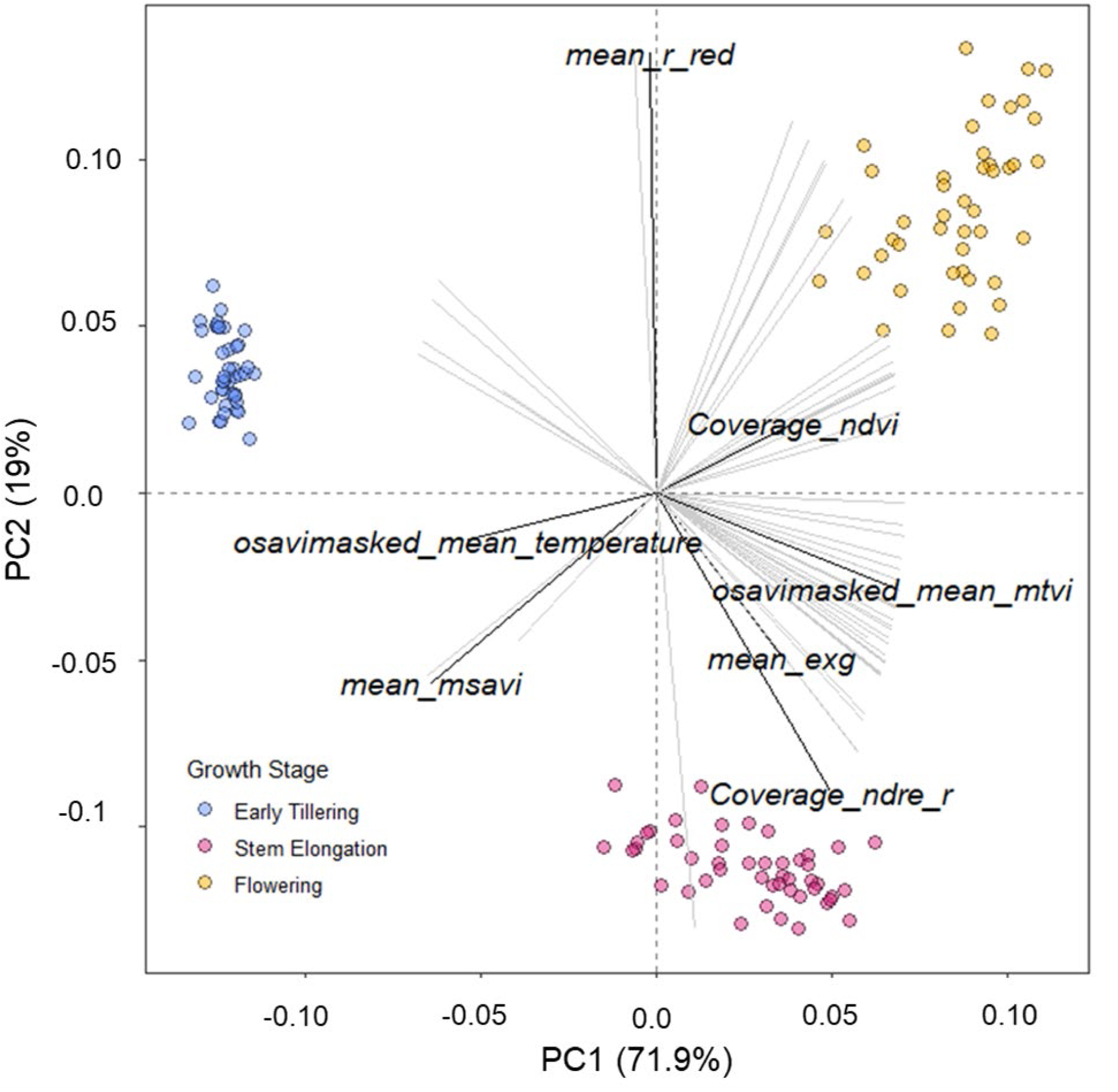
Biplot of vegetation indexes used to predict biomass at three growth stages. Principal component analysis (PCA) of various vegetation indexes is used to predict biomass at three distinct growth stages: Early Tillering, Stem Elongation, and Flowering. Seven vegetation indexes used in the final PLSR model are highlighted, and the vectors indicate the direction and magnitude of the influence of each vegetation index on their contribution to the variance in biomass prediction at different growth stages.

**Fig. S3.**
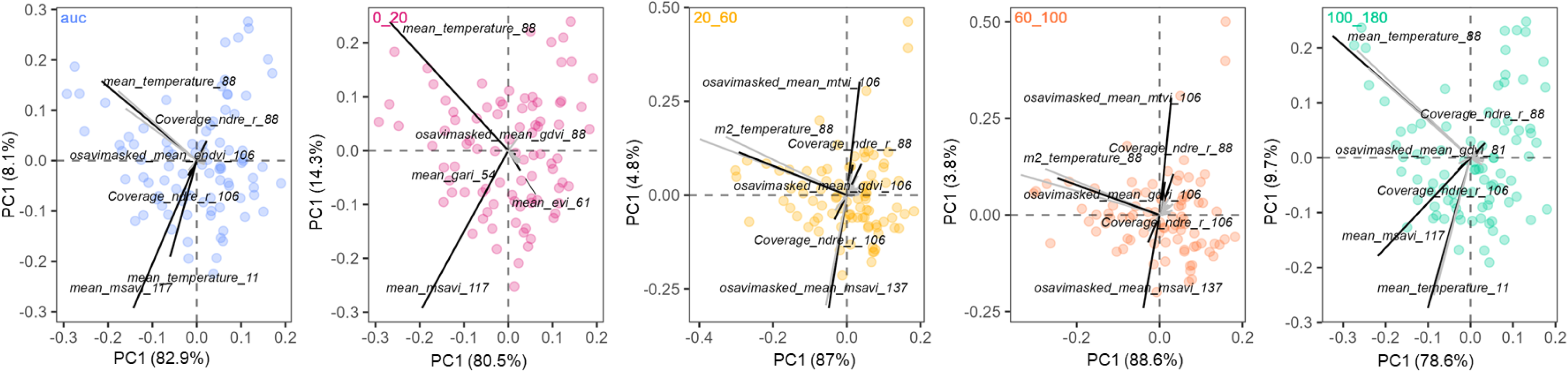
Biplot of vegetation indexes used to predict biomass at various root growth count curves. The set of biplots illustrates the principal component analysis (PCA) of the final vegetation indexes used to predict the area under the root growth count curve (auc) overall and across four different sections. Each biplot represents (left to right) auc (overall), 0-20, 20-60, 60-100, 100-180.

**Fig. S4.**
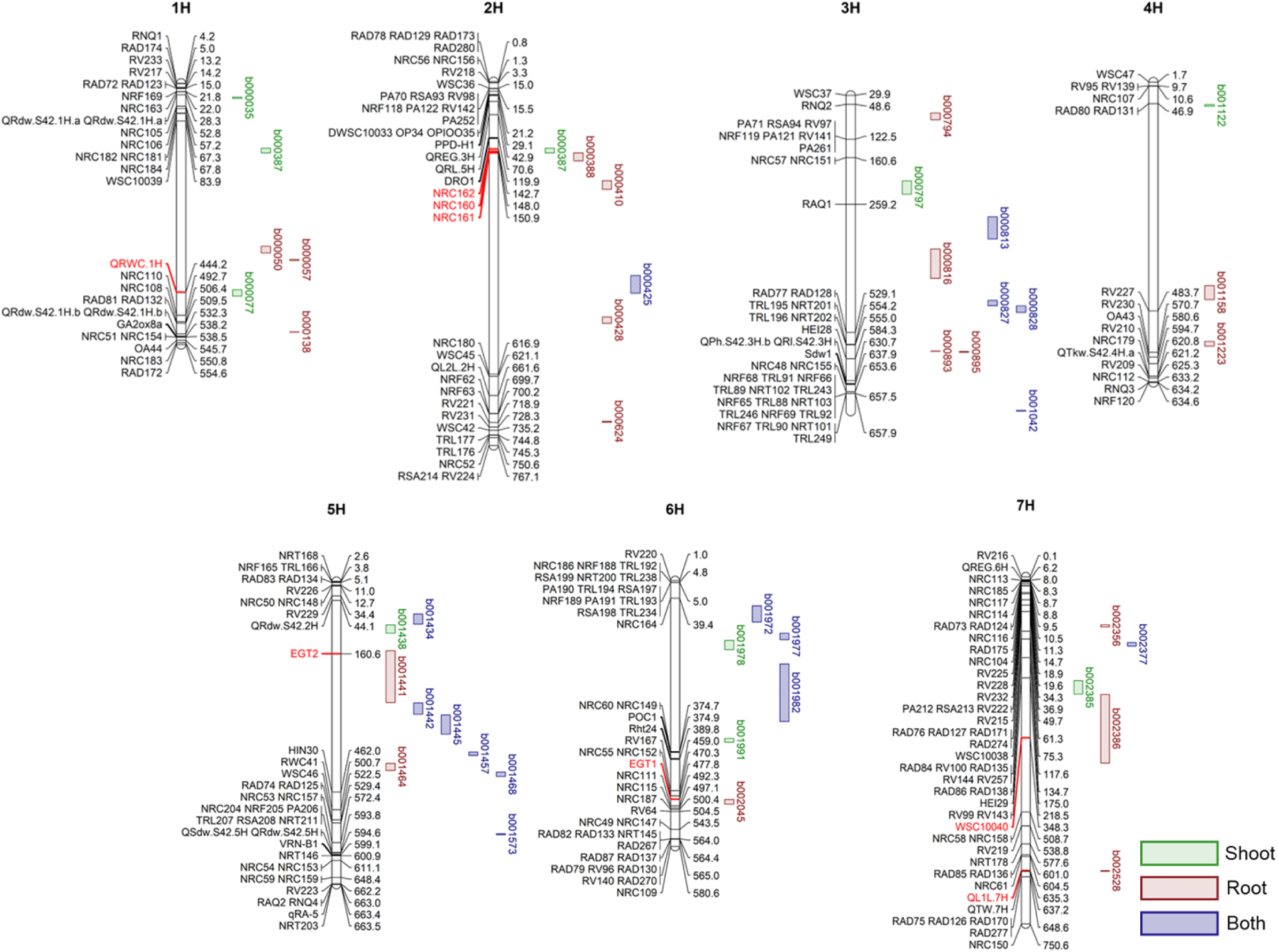
Alignment of haploblocks with barley RSA trait and canopy development QTLs across seven chromosomes (1H-7H). Previously reported QTLs on left of chromosomes are presented alongside physical positions Mbp to the right. Green highlights indicate novel significant haploblocks for canopy development (shoot). Brown indicates unique RSA traits haploblocks and blue shows overlap between RSA and shoot haploblocks identified in this study. Red markers represent alignment between previously reported QTL and haploblocks in this study.

**Table S1.** Calculated spectral vegetation indices and temperature-related variables are ranked based on correlation with dry biomass.

**Table S2.** Summary of predicted RSA traits for all plots, including minimum, maximum, mean and median values for overall root number and specific soil depth intervals (0–20 cm, 20–60 cm, 60–100 cm, and 100–180 cm)

**Table S3.** List of identified haplo-blocks in Australian barley breeding lines, their classifications, and associated predicted traits. Black indicates unique blocks, while red denotes blocks common across multiple traits.

**Table S4.** Previously reported QTLs and developmental genes in barley were used to align with identified haplo-blocks. Details of populations used, chromosomal locations, environmental contexts and references are provided.

